# Alkaline intracellular pH activates AMPK-mTORC2 signaling to promote cell survival during growth factor limitation

**DOI:** 10.1101/2021.05.13.444090

**Authors:** D Kazyken, SI Lentz, DC Fingar

## Abstract

mTORC2 controls cell metabolism and promotes cell survival, yet its upstream regulation by diverse cellular cues remains poorly defined. While considerable evidence indicates that mTORC1 but not mTORC2 responds dynamically to amino acid levels, several studies reported activation of mTORC2 signaling by amino acids, a paradox that remains unresolved. Following amino acid starvation, we noted that addition of a commercial amino acid solution but not re-feeding with DMEM containing amino acids increased mTORC2 signaling. Interestingly, the pH of the amino acid solution was ∼ 10. These key observations enabled us to discover that alkaline intracellular pH (pHi) represents a previously unknown activator of mTORC2. Using a fluorescent pH-sensitive dye (cSNARF-1-AM) coupled to live-cell imaging, we demonstrate that alkaline extracellular pH (pHe) increases intracellular pHi, which increases mTORC2 catalytic activity and downstream signaling to Akt. Alkaline pHi also activates AMPK, a sensor of energetic stress. Functionally, alkaline pHi attenuates apoptosis caused by growth factor withdrawal, which requires AMPK in part and mTOR in full. Collectively, these findings reveal that alkaline pHi increases AMPK-mTORC2 signaling to promote cell survival during growth factor limitation. As elevated pHi represents an under-appreciated hallmark of cancer cells, alkaline pH sensing by AMPK-mTORC2 may contribute to tumorigenesis.

**One Sentence Summary:** Alkaline intracellular pH activates mTORC2

## Introduction

mTOR (the mechanistic target of rapamycin) comprises the catalytic core of two distinct multi-protein complexes, mTORC1 and mTORC2. These mTORCs sense and integrate diverse extra- and intra-cellular signals derived from hormones, growth factors, nutrients, and energy through distinct downstream substrates to control cell physiology in ways appropriate for biological context (1-5). mTORC1 primarily promotes anabolic cellular processes (e.g., protein, lipid, and nucleotide synthesis) that sustain cell growth and cell proliferation while mTORC2 controls cell metabolism, cell survival, and the actin cytoskeleton (1-5). Not surprisingly, elevated mTORC1 and mTORC2 signaling contribute to pathologic conditions, including tumorigenesis (4,5).

Activation of mTORC1 by the cooperative action of insulin and amino acids has been studied extensively. In response to insulin, activation of the PI3K-Akt-TSC pathway leads to Rheb-GTP mediated activation of mTORC1 on the surface of lysosomes in a manner that requires sufficient levels of amino acids (4,6-8). Amino acids load RagA/B proteins with GTP, which recruit mTORC1 to lysosomal membranes in proximity to Rheb (8-14). Through an induced proximity mechanism, Rheb-GTP in turn interacts with and activates mTORC1 through conformational changes (15-17). TSC and Rheb represent central signaling nodes at which growth factor and amino acid signals converge to effect mTORC1 regulation (4,6,7). Insulin-PI3K-Akt signaling and amino acid sufficiency are required simultaneously to dissociate TSC from lysosomal membranes and away from Rheb, thus maintaining Rheb-GTP loading (18-20). Our knowledge of upstream pathways and mechanisms controlling mTORC2 activity and downstream signaling lags far behind that of mTORC1, however (4,5). Activation of mTORC2 signaling by hormones and growth factors requires PI3K (4,5,21), and oncogenic Ras interacts with mTORC2 to increase its activity on the plasma membrane (22,23). Curiously, nutrient withdrawal, specifically glutamine or glucose, activates mTORC2 (24,25), and upregulation of the stress sensing protein Sestrin2, which occurs during glutamine deprivation, increases mTORC2 activity (26,27). In addition, our prior work demonstrated that during energetic stress, AMP-activated protein kinase (AMPK) directly activates mTORC2 to promote cell survival (25).

Considerable evidence indicates that amino acids are required for mTORC1 but not mTORC2 signaling (4,5,28-31). Several studies reported paradoxical activation of mTORC2 signaling by amino acid stimulation, however (32-35). The reason for this discrepancy remains unclear. While studying amino acid sensing by mTORC1, we noted that after amino acid starving cells, addition of a commercial amino acid solution but not re-feeding cells with DMEM containing amino acids activated mTORC2 signaling. Interestingly, we found the pH of the amino acid solution to be ∼ pH 10. When we adjusted the pH of the amino acid solution to physiological pH 7.4, it failed to increase mTORC2 signaling. These key observations enabled us to discover and demonstrate here that alkaline extracellular pH (pHe) increases intracellular pH (pHi), which increases AMPK and mTORC2 signaling to attenuate apoptosis caused by growth factor withdrawal. As elevated pHi represents an under-appreciated hallmark of cancer cells (36-38), alkaline pH sensing by AMPK-mTORC2 may enable growth factor- and nutrient-deprived cancer cells at the core of a growing tumor to evade apoptosis and survive.

## Results

### Amino acids at alkaline pH but not physiological pH increase mTORC2 and AMPK signaling

Researchers in the mTOR field employ diverse methods to amino acid starve and stimulate cells. While studying the mTORC1 response to amino acids, we noted that after amino acid starving cells in D-PBS containing dialyzed FBS (dFBS), addition of a commercial amino acid solution increased mTORC2 signaling, as measured by the sensitivity of Akt S473 phosphorylation to the mTOR inhibitor Torin1 (Figure 1A, left). Consistent with Akt S473 phosphorylation promoting and/or stabilizing Akt T308 phosphorylation (25,39,40), the amino acid solution also increased Torin1-sensitive Akt T308 phosphorylation (Figure 1A, left). We obtained similar results in the absence of serum growth factors (i.e., without FBS) (Figure 1A, right). Curiously, amino acid stimulation also increased phosphorylation of AMPK*α* on its activation loop site (S172) as well as the phosphorylation of mTOR S1261 and ACC S79, direct substrates of AMPK (Figure 1A, left and right) (25,41,42). As expected, amino acids increased S6K1 T389 phosphorylation in a Torin1-sensitive manner, an established readout of mTORC1 signaling (Figure 1A). On their face, these data are consistent with prior reports that amino acids increase mTORC2 (32-35) and AMPK signaling (33).

**Figure 1:**
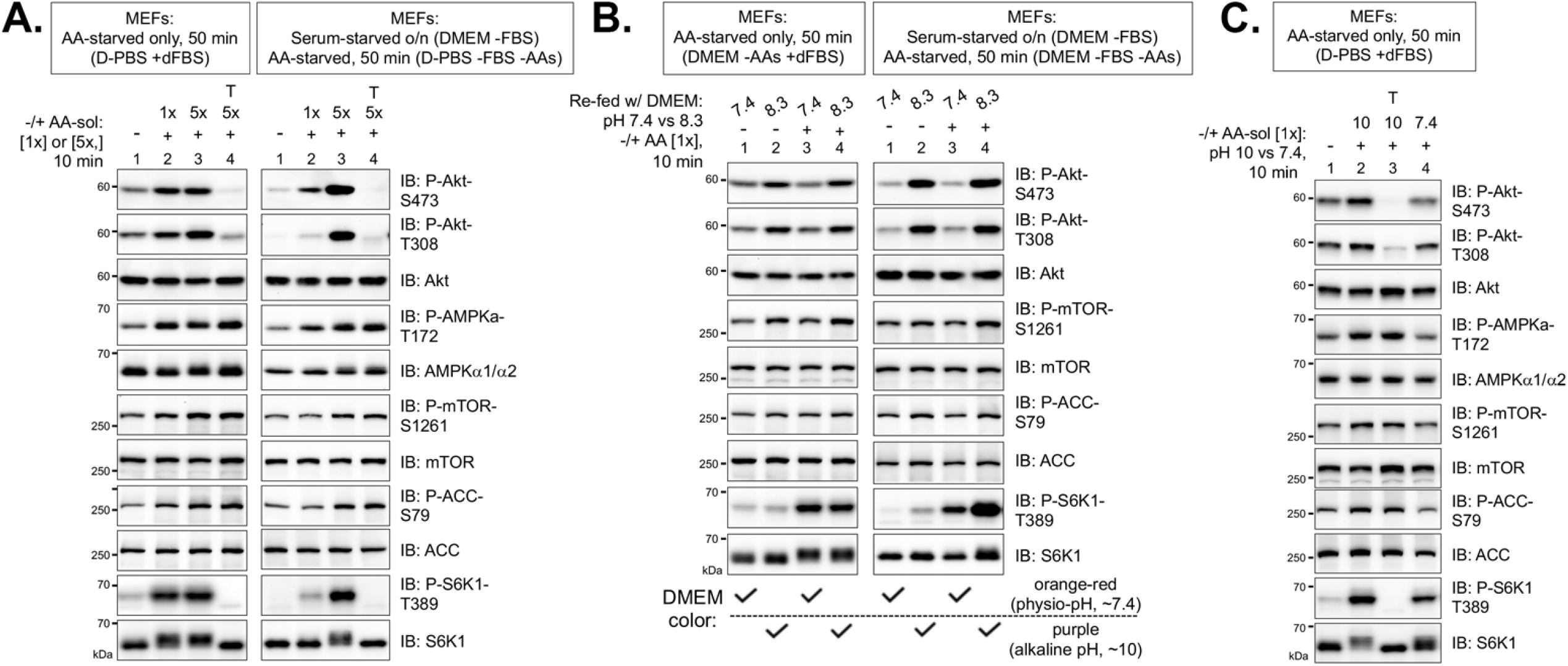
Amino acid stimulation at alkaline but not physiological pH increases mTORC2 and AMPK signaling. **A**. MEFs were cultured in complete media (DMEM/ +FBS) (left) or serum starved overnight, ∼16 hr. (right). They were next amino acid (AA) starved in D-PBS (50 min) with (left) or without (right) dFBS, pre-treated with Torin1 (T), and treated without (-) or with (+) a commercial amino acid solution (AA-sol) (10 min) to [1x] or [5x] final. Whole cell lysates were immunoblotted as indicated. **B**. MEFs were cultured as in **A** but amino acid (AA) starved in AA-free DMEM and re-fed with DMEM at pH 7.3 or 8.3 lacking (-) or containing (+) AAs (10 min). **C**. MEFs were amino acid starved in D-PBS/ dFBS, pre-treated with Torin1 (T), and stimulated with an AA solution at pH 10 or 7.4 (10 min).

When we used DMEM to amino acid deprive cells rather than D-PBS, however, addition of the amino acid solution induced a rapid change in DMEM color from orange-red (indicates physiological pH ∼7.4) to purple (indicates alkaline pH). Indeed, we measured the pH of the amino acid solution and found it to be ∼pH 10. To carefully compare how alkaline pH vs. amino acids control mTORC2 signaling, we adjusted the pH of DMEM lacking or containing amino acids to 7.4 or 8.3. DMEM at pH 8.3 but not 7.4 increased mTORC2 (i.e., Akt P-S473 and P-T308) and AMPK (i.e., mTOR P-S1261; ACC P-S79) signaling regardless of amino acid status in both the presence (Figure 1B, left) and absence (Figure 1B, right) of serum growth factors. As expected, DMEM containing but not lacking amino acids increased mTORC1 signaling (i.e., S6K1 P-T389) regardless of pH (Figure 1B). Upon adjusting the pH of the amino acid solution, its addition at pH 10 but not 7.4 to cells in D-PBS (+ dFBS) increased mTORC2 and AMPK signaling (Figure 1C). Importantly, our findings confirm those of Tato *et al*. (32) and Pezze *et al*. (33), who reported that commercial amino acid solutions added to amino acid starved cells increase mTORC2 signaling. Figure 1A (right) replicates the results of Tato *et al*. using identical amino acid starvation and stimulation conditions (Figure 1A right), while Figure S1A replicates the results of Pezze *et al*. in which addition of an amino acid solution to C2C12 myoblasts incubated in HBSS increased mTORC2 signaling. When we controlled for pH, however, addition of the amino acid solution at pH 7.4 to C2C12 myoblasts amino acid starved in either HBSS (Figure S1A) or DMEM (Figure S1B) failed to increase mTORC2 signaling. Collectively, these results provide strong evidence that increased pH-not amino acids-increases mTORC2 and AMPK signaling.

### Alkaline extracellular pH (pHi) increases mTORC2 catalytic activity and signaling

We next investigated whether alkaline extracellular pH (pHe) is sufficient to increase mTORC2 signaling in the absence of any changes in amino acid levels. Re-feeding MEFs cultured in complete media (DMEM with amino acids and FBS) with DMEM at pH 8.3 but not pH 7.4 increased Akt phosphorylation (P-S473 and P-T308) in MEFs (Figure 2A) and HEK293T cells (Figure S2A). These effects were rapid and transient, with maximal phosphorylation of Akt occurring at 5-15 minutes with apparent declines by 30-60 min, possibly because elevated pH cannot be maintained in these media conditions. As in Figure 1, the activating effect of pHe on Akt phosphorylation was Torin1-sensitive in MEFs (Figure 2B) and HEK293T cells (Figure S2B), thus reflecting mTORC2 signaling. pH dose response experiments (pH 7.4-9.0) demonstrated that pH 8.3-8.5 mediated maximal mTORC2 signaling in MEFs (Figure S2C). To test whether alkaline pHe increases mTORC2 intrinsic catalytic activity, we performed mTORC2 *in vitro* kinase (IVK) assays. We re-fed MEFs with DMEM pH 8.3, immuno-purified mTORC2 by immunoprecipitation of Rictor (a partner protein exclusive to mTORC2), and measured phosphorylation of His-Akt1 by mTORC2 *in vitro*. Alkaline pHe increased mTORC2 catalytic activity (Figure 2C). To increase pHi by an alternate method, we used NH_4_Cl, which increases cytosolic and organellar pH. Indeed, NH_4_Cl increased mTORC2 signaling (Figure 2D). Finally, we used the pharmacologic drug cariporide to lower intracellular pH (pHi), as in other studies (43-45). Cariporide inhibits NHEI, a H^+^-Na^+^ antiporter on the plasma membrane that drives proton (H^+^) efflux from the cytosol to the extracellular space (43-45). Cariporide blunted the ability of alkaline pHe to increase mTORC2 signaling in MEFs (Figure 2E) and HEK293 cells (Figure S1B). Taken together, these results demonstrate that increased intracellular (pHi) mediates the effect of alkaline extracellular (pHe) on mTORC2 signaling.

**Figure 2:**
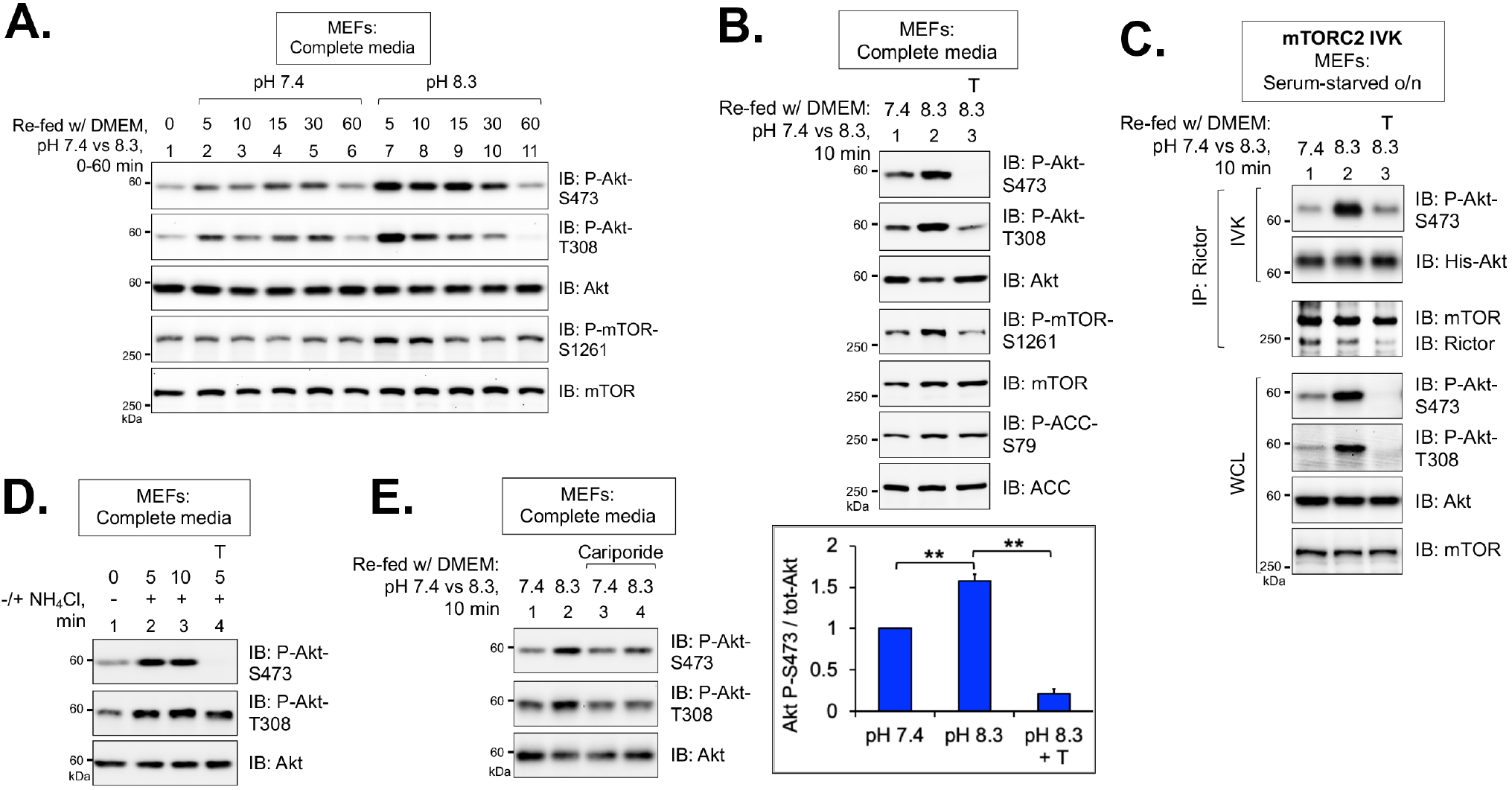
Alkaline extracellular pH (pHe) is sufficient to increase mTORC2 catalytic activity and signaling. **A**. MEFs in complete media (DMEM/ +FBS) were re-fed with media at pH 7.4 or 8.3 for various times (5-60 min). Whole cell lysates were immunoblotted as indicated. **B**. Similar to **A**, except cells were pre-treated with Torin1 (T). Graph, mean ratio ± SD of Akt P-S473/ Akt; n= 4 independent experiments. \*\**p* < 0.01 using one-way ANOVA and Tukey’s post hoc tests. **C**. Rictor was immunoprecipitated (IP) from MEFs that had been serum starved, pre-treated with Torin1 (T), and re-fed with serum-free DMEM at pH 7.4 or 8.3 (10 min) -/+ Torin1. *In vitro* kinase (IVK) reactions were performed with ATP and His-Akt1 substrate, with Torin1 present in the IVK reaction (lane 3). IVKs and WCLs were immunoblotted as indicated. **D**. MEFs in complete media (DMEM/ FBS) were pre-treated with Torin1 (T) and stimulated without (-) or with (+) NH_4_Cl (5-10 min.). **E**. Similar to **A**, except cells were pre-treated with cariporide (30 min).

### Incubation of cells in media at alkaline pH or containing NH4Cl increases intracellular pH (pHi)

We next confirmed that incubation of cells in media at alkaline pH increases intracellular pH (pHi). To do so, we used live-cell imaging coupled with a cell-permeable, pH-sensitive fluorescent dye (cSNARF-1-AM) that is capable of measuring changes in intracellular pH (pHi) between 7 and 8. cSNARF-1-AM undergoes a pH-sensitive shift in fluorescence wavelength emission depending on protonation state. With 488 nm excitation, peak emission occurs at 580 nm in the protonated state (more acidic) and 640 nm in the de-protonated state (more alkaline). Thus, ratiometric imaging (640:580 nm) enables detection of changes in intracellular pH within the physiological range of normal cells (∼pH 7.2) and cancer cells (∼ pH 7.4-7.6) (37,46). MEFs were pre-loaded with cSNARF-1-AM and then re-fed with complete DMEM at pH 7.4 for 10 minutes followed by re-feeding with media at pH 8.3 for 10 minutes. Visualization of the acquired pseudo-colored ratiometric images (640:580 nm) and quantitation of the signal ratios revealed a striking and significant increase in intracellular pH (**Figure 3A**). Addition of NH_4_Cl to DMEM (pH 7.4) also increased pHi (**Figure 3B**. These results confirm that alkaline extracellular increases intracellular pH, consistent with other studies (47,48).

**Figure 3:**
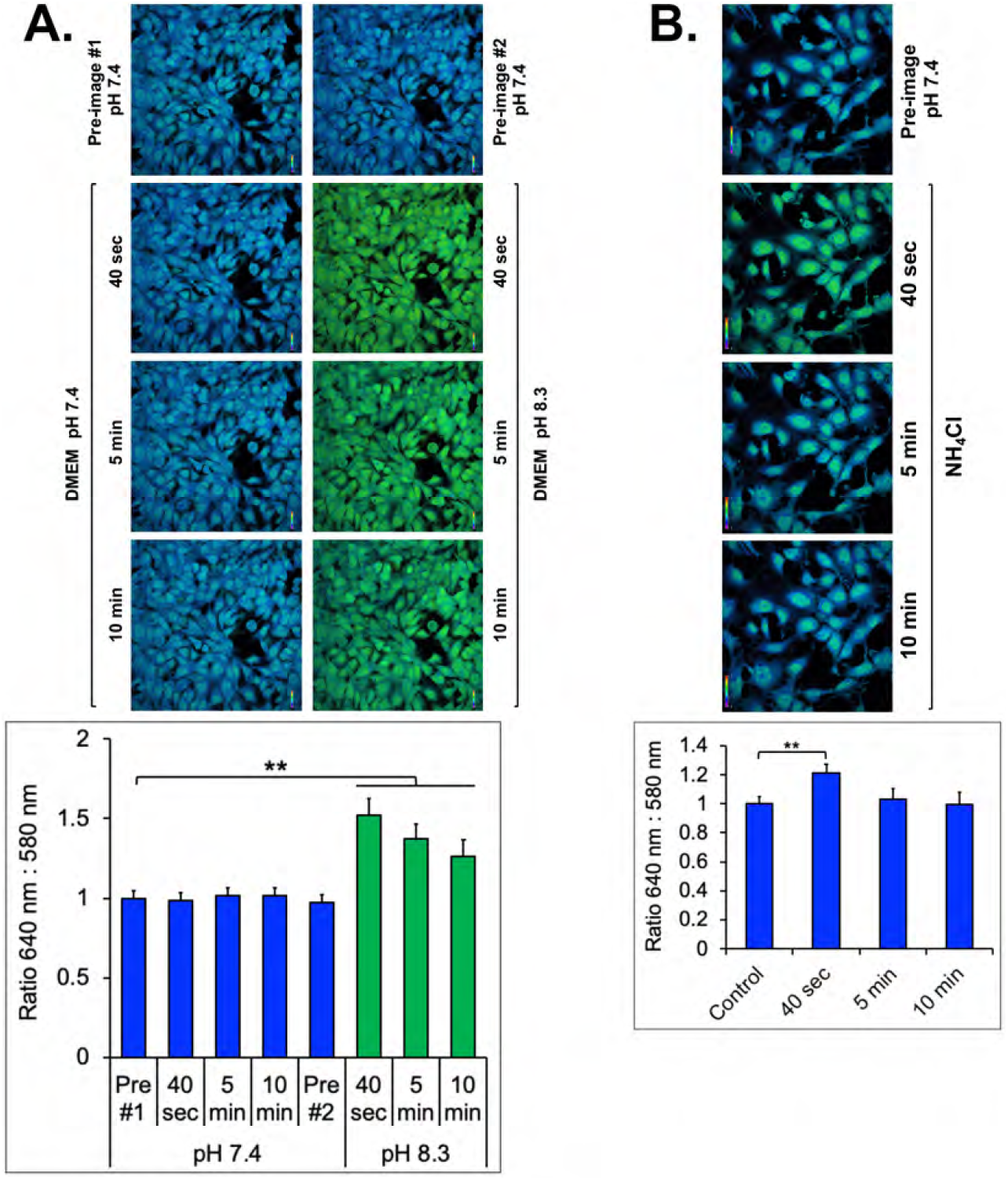
Incubation of cells in media at alkaline pH or containing NH4Cl increases intracellular pH (pHi) **A**. MEFs were pre-loaded with cSNARF-1-AM in in serum-free DMEM pH 7.4 (30 min). They were re-fed with DMEM/ FBS pH 7.4 and one image set was acquired (pre-image #1 pH 7.4). The cells were re-fed again with the same media, and three image sets were acquired at 40 sec., 5 min., and 10 min. At this point, the cells were re-fed once more with DMEM/ FBS at pH 7.4, and another image set was acquired (pre-image #2 pH 7.4). Next, the cells were re-fed with DMEM pH 8.3, and three image sets were acquired at 40 sec., 5 min., and 10 min. Ratiometric (580:640 nm) pseudo-colored images are shown for each treatment condition. Scale bars in the ratiometric images are pseudo-colored between values of 0 (violet) and 2.0 (red). Graph, quantitation of ratiometric images, n=100 cells from 4 fields (∼25 cells/field) +/- SD. **p<.01 using one-way ANOVA and Tukey’s post hoc tests. **B**. MEFs were loaded with cSNARF-1-AM and treated as in **A**, except image sets were acquired from cells incubated in DMEM /FBS pH 7.4 without (pre-image) or with N_H_4Cl for 40 sec., 5 min., or 10 min. Ratiometric (580:640 nm) pseudo-colored images are shown for each treatment condition. Graph, same as above. **p<.01 using ANOVA as above.

### AMPK promotes mTORC2 signaling and cell survival in response to alkaline intracellular pH (pHi)

Our prior work demonstrated that energetic stress increases mTORC2 catalytic activity and signaling directly through AMPK mediated phosphorylation of mTOR and mTORC2 partner proteins (e.g., Rictor) (25). As the AMPK-mTORC2 axis responds to energetic stress, we speculated that it may also respond to alkaline pH stress. We therefore compared the ability of alkaline pHe to increase mTORC2 signaling in wild type and AMPK*α*1/*α*2 double knockout MEFs (i.e., AMPK DKO). When MEFs incubated in D-PBS/ dFBS were stimulated with the amino acid solution at pH 10 or re-fed with D-PBS/ dFBS adjusted to pH 9.5 (the pH of D-PBS/ dFBS after addition of the pH 10 amino acid solution to 1x), we found that alkaline pHe increased mTORC2 signaling in a manner partly dependent on AMPK (Figures 4A, 4B). As expected, AMPK double knockout abrogated AMPK signaling (AMPK P-S172; mTOR P-S1261; Raptor P-S792) (Figures 4A, 4B). Alkaline pHi was also sufficient to increase mTORC2 signaling in a manner partly dependent on AMPK in cells cultured in complete media (i.e., DMEM/ FBS) (Figure 4C). These results indicate that alkaline pHi increases mTORC2 signaling through AMPK and another unknown signal(s).

**Figure 4:**
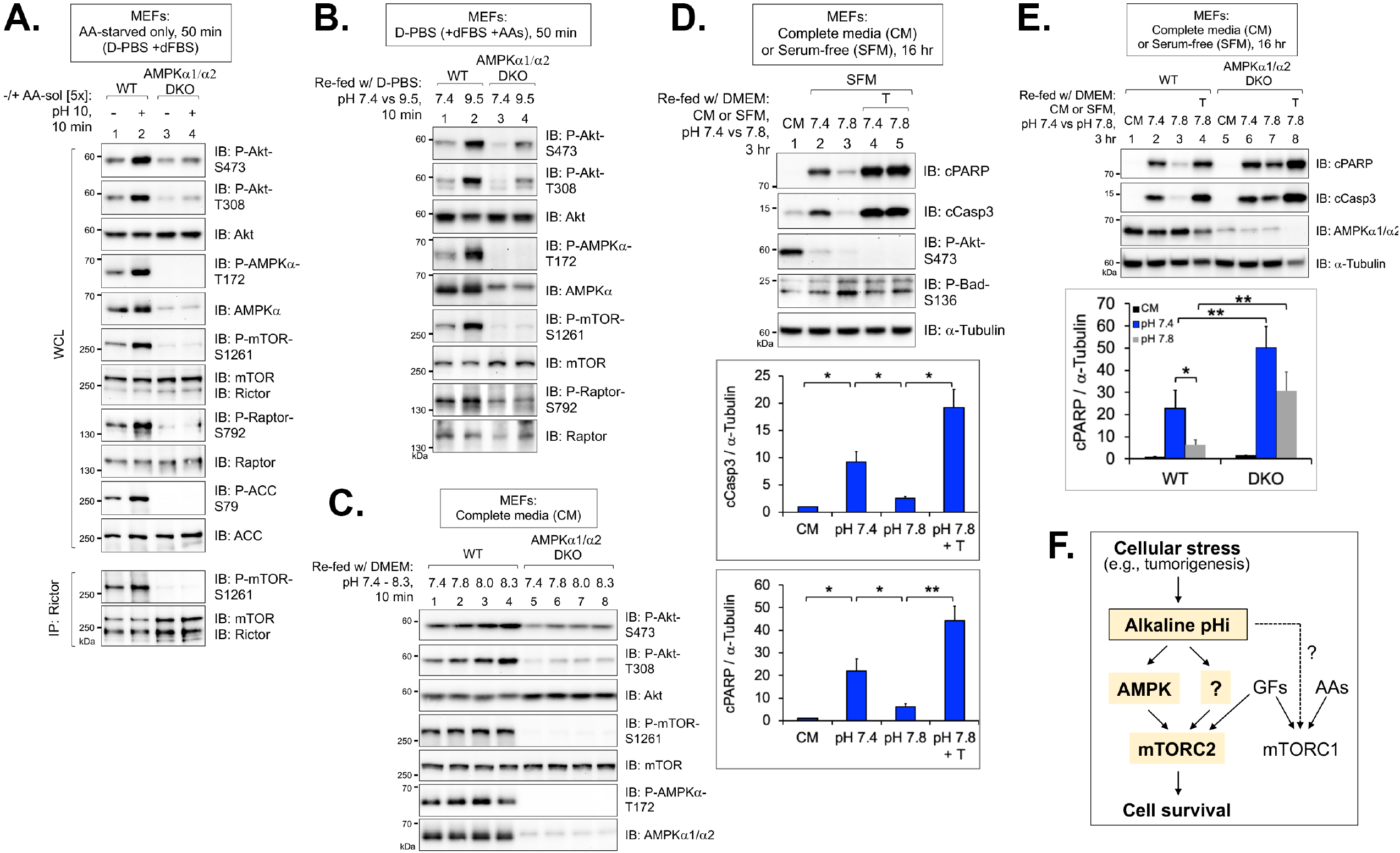
AMPK promotes mTORC2 signaling and cell survival in response to alkaline intracellular pH (pHi) **A**. Wild type (WT) and AMPK*α*1/*α*2 double knockout (DKO) MEFs were amino acid starved in D-PBS/ +dFBS (50 min) and stimulated with an amino acid solution whose pH had not been adjusted (i.e., pH=10) to [5x] final (10 min). Rictor was immunoprecipitated (IP), and IPs and whole cell lysates (WCL) were immunoblotted as indicated. **B**. WT and AMPK DKO MEFs were cultured in D-PBS/ +dFBS containing 1x amino acids (AAs) and re-fed with the same media at pH 7.4 or 9.5 (10 min). Note that pH 9.5 is the pH of D-PBS/ +dFBS supplemented with pH 10 AAs to 1x final. **C**. WT and AMPK DKO MEFs in complete media (DMEM/ +FBS) were re-fed with the same media at various pH values, 7.4-8.3 (10 min). **D**. WT MEFs cultured in DMEM complete media (CM) (DMEM/ +FBS) or serum-free media (SFM) for 16 hr were re-fed with CM at pH 7.4 or SFM pH 7.4 or 7.8 without or with Torin1 (T) for an additional 3 hr. (19 hrs. total). Graph, mean ratio ± SD of cParp / tubulin and cCasp3/ tubulin. n = 4 experiments. \**p* < 0.05, ** *p* < 0.01 using one-way ANOVA and Tukey’s post hoc tests. **E**. WT and AMPK*α*1/*α*2 DKO MEFs were treated as in **D**. Graph, mean ratio ± SD of cParp / tubulin. n = 5 experiments. \**p* < 0.05, ** *p* < 0.01 using ANOVA as above.

Cell survival requires sufficient levels of growth factors (49), and our recent work demonstrated that AMPK-mTORC2 signaling promotes cell survival in response to acute energetic stress (25). We therefore tested the hypothesis that elevated pHi protects against apoptosis during growth factor limitation, a setting common for growing tumors. We therefore serum-starved MEFs overnight (16 hr.) in DMEM and then re-fed the cells with either complete media (i.e., with FBS) or serum-free media (i.e., without FBS) for an additional 3 hrs. As expected, cells maintained in serum-free media for the full 19 hrs. displayed increased apoptosis relative to those rescued with complete media for the last 3 hrs., as monitored by blotting for cleaved caspase 3 (cCas3) and cleaved Parp (cParp) (Figure 4D). Consistent with our hypothesis that alkaline pHi protects against apoptosis, re-feeding MEFs with serum-free media whose pH had been adjusted to 7.8 using sodium bicarbonate rather than NaOH (which maintains the pH of DMEM for longer periods of time) suppressed apoptosis relative to serum-free re-feeding at pH 7.4 in a Torin1 sensitive manner (Figure 4D). While prolonged alkaline pHi (i.e., 3 hr.) could not maintain Akt S473 phosphorylation, it maintained inhibitory phosphorylation of the Akt substrate and pro-apoptotic protein BAD (Figure 4D). Consistent with our hypothesis that AMPK is required for alkaline pHi to protect against apoptosis, double knockout of AMPK*α*1/*α*2 partially suppressed the ability of serum-free media at alkaline pH 7.8 to attenuate apoptosis caused by growth factor withdrawal (Figures 4E; S3).

## Discussion

This study identifies alkaline intracellular pH (pHi) as a previously unrecognized activator of AMPK and mTORC2 that attenuates apoptosis during growth factor limitation (**Figure 4F**). As AMPK promotes mTORC2 signaling and cell survival mediated by alkaline intracellular pH (pHi) in part rather than in full, other signals (?) cooperate with AMPK to activate mTORC2 in response to alkaline pHi. It is important to note that a recent study found that alkaline pHi increased Akt (S473) phosphorylation, although mechanistic details were not defined (47). We speculate that prior studies reporting activation of AMPK and mTORC2 signaling by amino acids (32-35) likely mistook an increase in pHi for an increase in amino acid levels. In fact, Tato *et al*. concluded that amino acids selectively activate mTORC2 signaling depending on the method of amino acid starvation and stimulation employed (32) (i.e., amino acids activated mTORC2 signaling in amino acid starved cells in response to addition of a commercial amino acid solution but not upon re-feeding with amino acid replete DMEM). By controlling for the pH of the amino acid solution or using DMEM lacking or containing amino acids, our results reveal that alkaline pHi-not amino acids-increases mTORC2 signaling, thus correcting a misconception in the mTOR field.

Dysregulated pH represents an under-appreciated hallmark of cancer cells, which display a reversal in the pH gradient controlled by altered proton (H^+^) flux (i.e., elevated intracellular pHi and decreased extracellular pHe, the reverse of normal cells) (36-38,45,46). Functionally, dysregulated pH dynamics modifies cancer cell behaviors, including proliferation, survival, metabolic adaptation, migration, and metastasis. Changes in pHi control the structure and function of pH-sensitive proteins (aka, pH-sensors) through protonation/ de-protonation, a posttranslational modification akin to phosphorylation, ubiquitination, etc. (36-38,46). Documented pH-sensors with recurring charge-changing mutations (e.g., Arg to His) include p53, EGF-receptor, Ras-GRP1, and *β*-catenin (50-53). Elevated activity of several plasma membrane ion exchangers, including the Na^+^-H^+^ exchanger NHE1, contributes to increased pHi and correlates with tumor initiation, progression, and metastasis (37,45). Indeed, increased H^+^ efflux contributes to transformed cell behaviors mediated by oncogenic Ras, as demonstrated in human mammary cells and a Drosophila model *in vivo* (54). By identifying alkaline pHi as an activator of AMPK-mTORC2 signaling that attenuates apoptosis caused by growth factor withdrawal, our work suggests that the AMPK-mTORC2 axis may drive tumorigenesis in response to dysregulated pH dynamics by enabling growth factor-, nutrient-, and oxygen-deprived cancer cells at the core of a growing tumor to survive. Such a role is consistent with the paradoxical role of AMPK as a tumor promoter in certain contexts (despite its established role as a tumor suppressor) and the newfound role for AMPK-mTORC2 signaling in cell survival during energetic stress (25,55-60). More broadly, our results suggest that the AMPK-mTORC2 axis senses diverse types of cellular stress, which likely rewires cell metabolism to help cells adapt and survive. In the future, it will be important to identify the pH-sensors that transduce signals to AMPK and mTORC2.

## Experimental Procedures

### Cell Culture

Cell lines (MEFs; HEK293T; C2C12) were cultured in DMEM containing high glucose [4.5 g/liter], glutamine [584 mg/liter] and sodium pyruvate [110 mg/liter] (Life Technologies #11995-065) supplemented with 10% fetal bovine serum (FBS) (Life Technologies #10347-028) and incubated at 37°C in a humidified atmosphere with 7.5% CO_2_. To effect amino acid (AA) starvation, cells were cultured in either D-PBS (which contains [1 g/L] D-glucose) (Life Technologies #14287-080) or AA-free DMEM (US Biologicals #D9800-13) without or with 10% dialyzed FBS (Life Technologies #A33820-01) for 50 minutes. To effect serum-starvation, cells were cultured in DMEM containing 20 mM Hepes pH 7.2 for ∼16 hr. overnight. Cells were amino acid (AA) stimulated in two ways: **1)** An amino acid solution (RPMI 1640 Amino Acid Solution [50x] (Sigma # R7131) supplemented with L-glutamine (Sigma # G8540) was added to cells in AA-free media to a final concentration of 1x (∼ concentration in RPMI) or 5x (∼ concentration in DMEM). The pH of the amino acid solution was either not adjusted (∼pH 10) or adjusted to pH 7.4 with 1N HCl; or **2)** Cells were re-fed with AA-replete DMEM after incubation in AA-free DMEM. To treat cells at various alkaline extracellular pHs acutely, the pH of D-PBS or DMEM was adjusted with NaOH. To stably maintain DMEM at alkaline pH (i.e., pH 7.8), the NaHCO_3_ concentration was increased from 44 mM to 120 mM by directly dissolving in NaHCO_3_ followed by filter sterilization and equilibration overnight in an incubator at 7.5% CO_2_, as described (48).

### Live-cell imaging of cSNARF-1-AM

MEFs were plated in 8-well cover glass bottom chambers (Lab-tek #155409) and loaded with cSNARF-1-AM [10 μM] (Thermo Fisher #C1272), a fluorescent pH-sensitive dye, for 30 min at 37°C in serum-free DMEM. The cells were then rinsed with serum-free DMEM and re-fed with DMEM/ FBS [10%] pH 7.4. Live-cell imaging (0-10 min) of cells cultured in DMEM/ FBS pH 7.4 or 8.3 was performed using a Nikon A1 confocal microscope equipped with a stage-top incubator that maintains temperature and CO_2._ Images were acquired using a 40x oil objective (1.3 NA) at a resolution of 1024×1024 and an optical thickness of 1.18 µm (confocal aperture set at 3 airy units). cSNARF1-AM signal was excited with an argon laser at 488 nm and images sets were collected simultaneously in two emission band pass filters at 553-618 nm (for 580 nm) and 663-738 nm (for 640 nm). Nikon Elements software was used to create and pseudo-color the 640:580 nm ratiometric images with values ranging from 0 (violet) to 2.0 (red). MetaMorph software was used to quantify the ratiometric images by measuring the average ratio of a region of interest within one cell, and 25 cells per image were quantified. A change in the 640:580 nm ratio and accompanying pseudo-colored image reflects a change in intracellular pH (pHi).

## Acknowledgments

We thank Drs. B. Viollet (Inserm, Paris, France) and R. Shaw (Salk Institute, San Diego, CA) for sharing immortalized wild-type and AMPK*α*1/*α*2 DKO MEFs. We also thank Dr. P. Goforth for advice with pH sensing dyes. This work was supported by a grant from the NIH (R01 GM-137577) (to DCF) and by the Michigan Diabetes Research Center (MDRC) (NIH-NIDDK; #P30 DK020572).

## Supplemental Information

See the Supplement for additional **a)** Experimental Procedures including materials, antibodies, drug treatments, cell lysis, immunoprecipitation, and western blotting, mTORC2 *in vitro* kinase assays, statistical analysis, and image editing; and **b)** Supporting Figures S1-S3.

## Abbreviations not defined in the text

mTORC1: mTOR complex 1
mTORC2: mTOR complex 2
mTORC: mTOR complex
D-PBS: Dulbecco’s-phosphate buffered saline
DMEM: Dulbecco’s modified eagle medium
HBSS: Hank’s balanced salt solution
MEFs: mouse embryonic fibroblasts
PI3K: phosphatidylinositol 3-kinase
NH_4_Cl: ammonium chloride

## Supplementary Information Supplemental Experimental Procedures

### Materials

General chemicals were from Thermo Fisher Scientific or Sigma Aldrich. NP40, Brij35, and CHAPS detergents were from Pierce; cOmplete Protease Inhibitor Cocktail (EDTA-free) tablets were from Millipore Sigma (#11836170001); protein A-Sepharose CL-4B was from Sigma-Aldrich (#17-0780-01); Immobilon-P polyvinylidene difluoride (PVDF) membrane (0.45 μM) was from Millipore; reagents for enhanced chemiluminescence (ECL) were from Alkali Scientific (Bright Star #XR92) or Advansta (WesternBright Sirius HRP substrate); NH_4_Cl was from Fisher (#A661); PageRuler and PageRuler Plus prestained protein ladders were from Thermo Fisher (#26617; 26619).

### Antibodies

The following antibodies were from Cell Signaling Technology (CST): AMPK*α* P-T172 (#4188); pan-AMPK*α* (#2532); Akt P-S473 (#4060); Akt P-T308 (#4056); Akt (#9272); mTOR (#2972); raptor P-S792 (#2083); raptor (#2280); S6K1 P-T389 (#9234); ACC P-S79 (#3661); ACC (#3676); Bad P-S136 (#4366); cleaved caspase 3 (#9664); cleaved PARP (#9544); *α*-tubulin (#2144); mTOR P-S2481 was from Millipore (#09-343). The following custom polyclonal anti-peptide antibodies were generated by us with the aid of Covance, as described previously (61): mTOR P-S1261 (amino acids 1256-1266; rat); rictor (amino acids 6-20; human); and S6K1 (amino acids 485-502; rat 70 kDa isoform). Donkey anti-rabbit-HRP secondary antibody was from Jackson (#711-095-152).

### Drug treatments, cell lysis, immunoprecipitation, and western blotting

Cells were pre-treated with Torin1 [100 nM] (30 min.) (shared by D. Sabatini, MIT and Whitehead Institute, Boston, MA) or cariporide [10 μM] (30 min.) (Sigma #SML1360) prior to lysis. Unless otherwise indicated, cells were washed 2x in PBS and lysed in buffer containing NP-40 [0.5%] and Brij35 [0.1%], as described (61). Lysates were incubated on ice (15 min.) and then spun at 13,200 rpm (5 min.) at 4°C. Post-nuclear supernatants were normalized for protein levels by Bradford assay. For immunoprecipitation, whole cell lysates were incubated with antibodies for 2 hr. at 4°C and Protein A-Sepharose beads for 1 hr. Beads were washed three times in lysis buffer and resuspended in 1x sample buffer. Samples were resolved on SDS-PAGE and transferred to PVDF membranes in Towbin transfer buffer containing 0.02% SDS. Western blotting was performed by blocking PVDF membranes in Tris-buffered saline (TBS) pH 7.5 with 0.1% Tween-20 (TBST) containing 3% non-fat dry milk, as described, and incubating the membranes in TBST/ BSA (2%) containing primary or secondary HRP antibodies. Blots were developed by ECL and detected digitally with a Chemi-Doc-It System (UVP).

### mTORC2 *in vitro* kinase assays

mTORC2 in vitro kinase (IVK) assays were performed, as described (25,62). Briefly, serum starved MEFs were re-fed with DMEM pH 7.4 or pH 8.3 and lysed in buffer containing CHAPS [0.3%]. Rictor was immunoprecipitated from a near confluent 10 cm plate and incubated with ATP [250 µM] and recombinant His-Akt1 [100 ng/reaction] (Millipore #14-279) in 15 µl kinase buffer (25 mM HEPES; 100 mM potassium acetate; 1 mM MgCl_2_) at 30°C for 30 min. and stopped by addition of sample buffer followed by incubation at 95°C for 10 min.

### Image editing

Adobe Photoshop was used for preparation of western blot images, using only levels, brightness, and/or contrast equivalently over the entire image. All presented images reflect the raw images.

### Statistical analysis

Fiji was used to quantitate ECL western blot signals. Results are presented as mean ± SD. Significance of the difference between two measurements was determined by Student’s t test. Multiple comparisons were analyzed for significance using one-way ANOVA followed by pairwise Tukey’s post hoc tests. Values of *p* <0.05 were considered significant. All experiments were performed at least three times, if not more, unless indicated otherwise in the figure legend.

## Supplementary Figure Legends

**Supplementary Figure 1:**
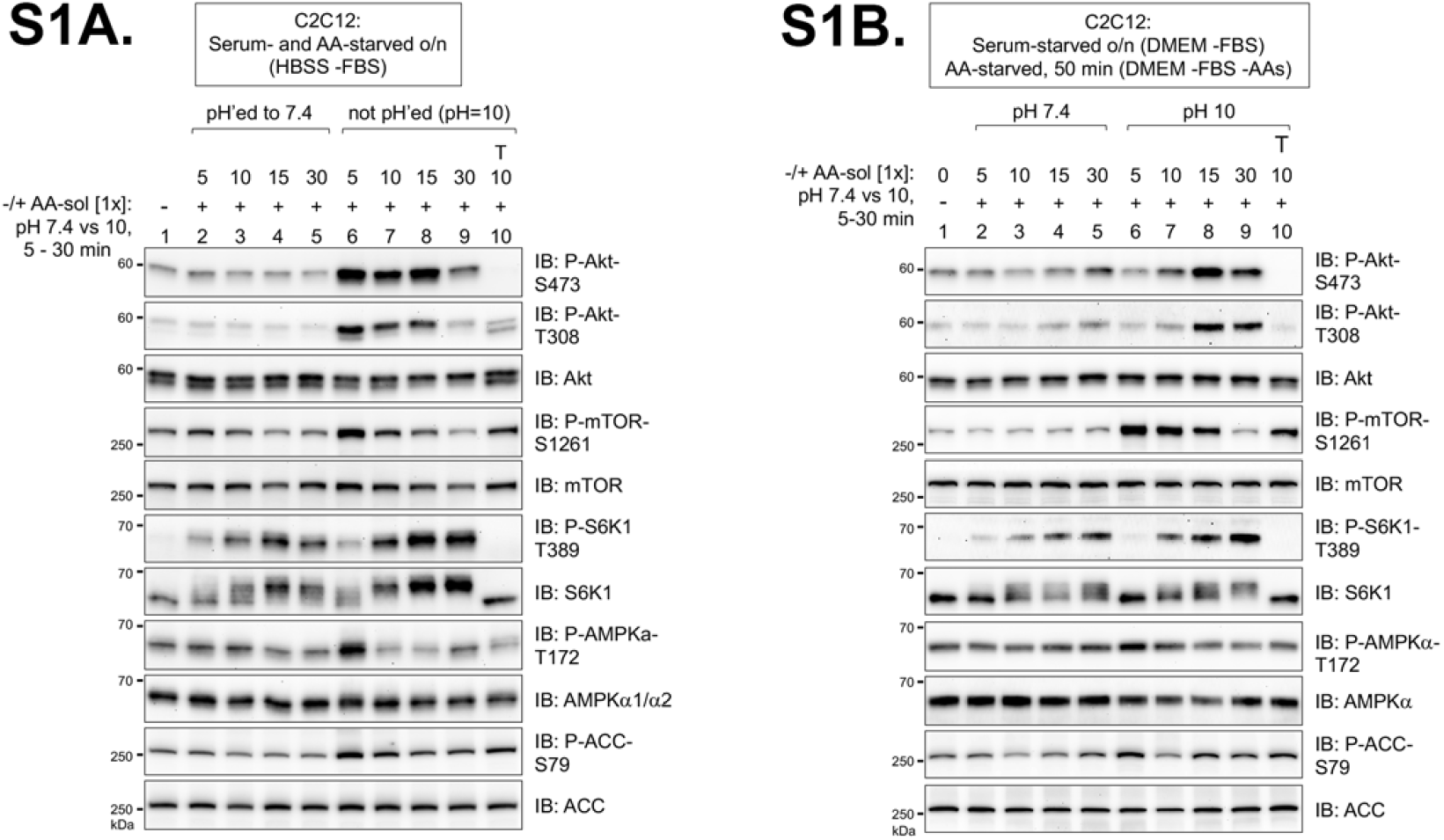
Activation of mTORC2 signaling by alkaline extracellular pH (pHe) in C2C12 myoblasts (related to figure 1) **S1A**. C2C12 cells were serum and amino acid starved overnight (∼16 hr) in HBSS and stimulated with an acid solution at pH 7.4 or 10 to 1x final (10 min). Whole cell lysates were immunoblotted as indicated. **S1B**. C2C12 cells were serum starved overnight in DMEM (∼16 hr.), shifted to amino acid-free DMEM lacking FBS (50 min), and then stimulated with an amino acid solution at pH 7.4 or 10 to 1x final (10 min).

**Supplementary Figure 2:**
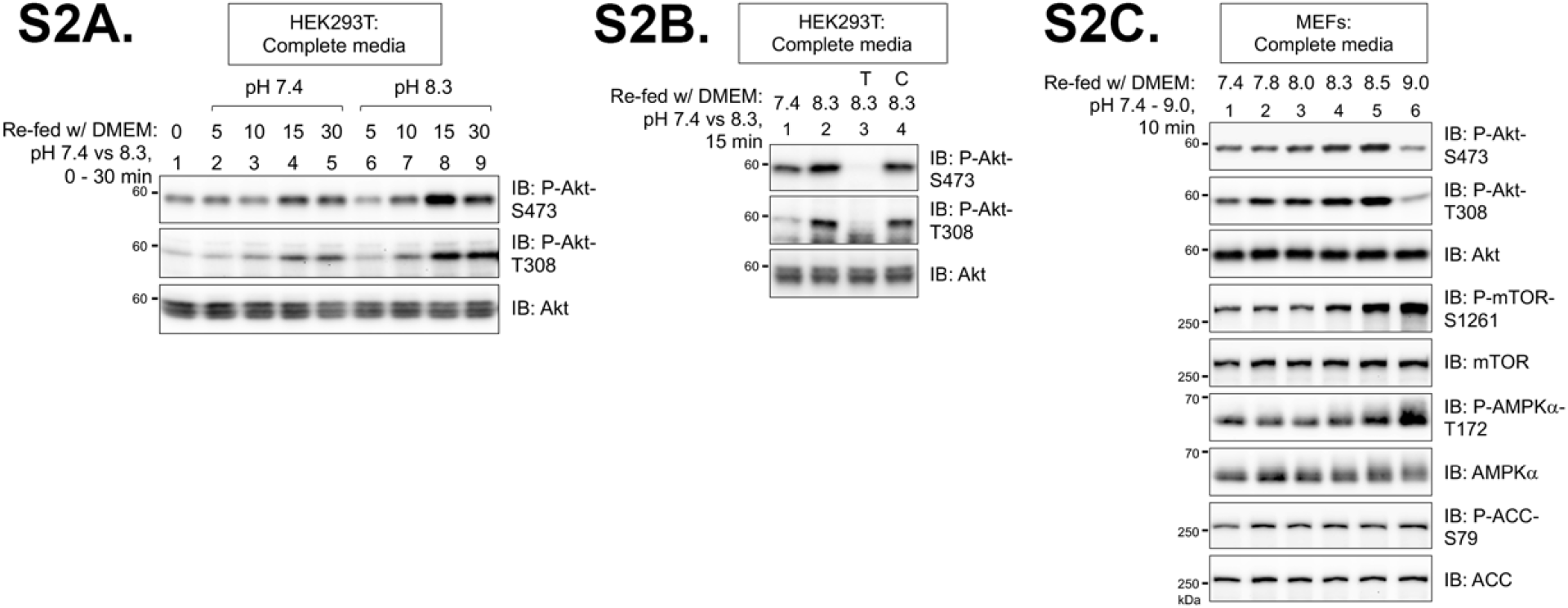
(related to figure 2) **S2A and S2B. Alkaline extracellular pH (pHe) increases mTORC2 signaling in HEK293T cells: S2A**, HEK293T cells were re-fed with complete media (DMEM/ FBS) (30 min) and then re-fed again with media at pH 7.4 or 8.3 for various times (0-60 min). Whole cell lysates were immunoblotted as indicated. **S2B**, HEK293T cells were pre-treated without or with Torin1 (T) or cariporide (C), and re-fed with complete media (DMEM/ FBS) at pH 7.4 or 8.3 (15 min) without or with drugs. **S2C. Alkaline pHe dose response in MEFs**. MEFs in complete media (DMEM/ FBS) were re-fed with media at various pH values (pH 7.4-9.0) (10 min).

**Supplementary Figure 3.**
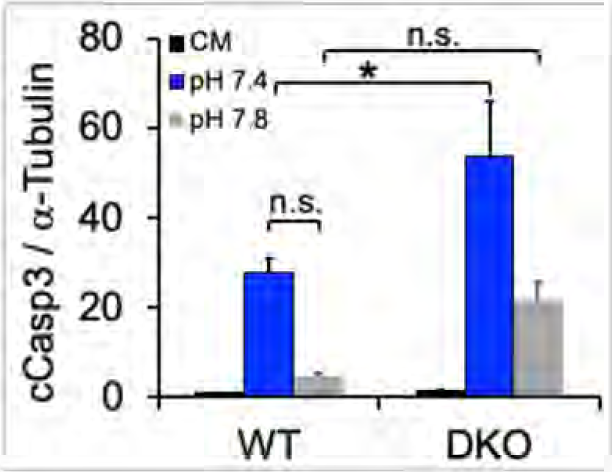
(related to figure 4)

## Notes

The authors declare that they have no conflicts of interest with the contents of this article.

